## Supplementary Results and figure supplemets for "Transcriptomic atlas of midbrain dopamine neurons uncovers differential vulnerability in a Parkinsonism lesion model"

**Figure 1-figure supplement 1**

**FANS plots for intact, lesioned, and untreated nuclei sorting**

Representative FACS plots, enriching for midbrain dopaminergic nuclei. PE-Texas Red-A (y-axis) represents mCherry channel. A-B) The DAPI$^{+}$Cherry$^{+}$ gates were set to select for all mCherry$^{+}$ nuclei in intact (A) and lesioned (B) samples. C-D) Control samples' gates were first set to select for all DAPI$^{+}$ nuclei (C), followed by gating for NeuN$^{+}$Cherry$^{+}$ nuclei (D). APC-A (x-axis) represents NeuN antibody channel.

**Figure 1-figure supplement 2**

**Quality control of snRNAseq data**

Metrics and indexes of data quality control before QC filtering (A-E). Distribution of genes (A) and UMIs (B) in libraries per condition. Cells' probability density plot for UMI/gene per condition (C). Correlation of UMIs to Genes, shown as a scatter plot for conditions (D). Percentages of mitochondrial and ribosomal transcripts (UMI) per condition (E). (F-J) The same metrics and indexes after QC filtering.

**Figure 1-figure supplement 3**

**Spatial mapping of main non-mDA-clusters**

A) A UMAP projection of all the cells with color-coding key. UMAP projections of *Slc17a7* (*Vglut1*, green clusters), *Slc17a6* (*Vglut2*, blue clusters) and *Gad2* (red and orange clusters) show the distribution of glutamatergic and GABAergic neuronal subtypes in various clusters. Cluster 3 contained a heterogenous mix of various midbrain and hypothalamic neurons expressing both glutamatergic, histaminergic and GABAergic markers. B) *Slc17a7*-expressing neuronal populations with example enriched genes, with cluster numbers and color-coding indicated. The *Slc17a7*$^{+}$ neurons were mainly found in cortex (CTX) and hippocampus, as well as in the pontine nuclei (Pn). The largest *Slc17a7*$^{+}$ cluster (2) contained mixed cortical neurons. B) *Slc17a6*$^{+}$ clusters were found in mostly in the midbrain and hypothalamus, for example in the medial geniculate nucleus (MGN), ventromedial hypothalamus (VMH), and retromammillary nucleus (RM). D) Serotonergic neurons expressing *Tph2* in dorsal raphe (DR). E) GABAergic clusters expressing *Gad2* were found dentate gyrus (DG), zona limitans intrathalamica (ZLI) and reticular nucleus (Rt). Cluster 8 co-expressed several DA genes, matching hypothalamic DA neurons. F) A dotplot representation of the enriched genes in non-mDA clusters.

**Figure 2-figure supplement 1**

**Distribution of *Th* mRNA and protein in different mDA populations**

(A-G) In situ hybridization and immunohistochemistry for *Th* mRNA and protein across the midbrain in 2 months old *Dat*$^{Cre/+}$*; R26*$^{TrapC/+}$ mice. In most of the RFP$^{+}$ cells both *Th* mRNA and protein can be detected. We detected some cells which were labelled with Dat-Cre and expressed mRNA but lacked TH protein (dotted circles), and some which expressed only *Th* mRNA but lacked both RFP and TH protein (dashed circles). These cells were not concentrated in any particular mDA neuron nuclei but were found scattered across VTA and SNpc. We also detected very rare cells in the medial regions of VTA and PAG which were labelled with Dat-Cre but lacked both *Th* mRNA and protein (solid circles). Aq, aqueduct; SNpc, Substantia Nigra pars compacta; PBP, Parabrachial Pigmented Nucleus; IF, Interfascicular Nucleus; PAG; Periaqueductal Grey. Scalebars 100 μm.

**Figure 3-figure supplement 1**

**Territory and neighborhood markers of mDA dataset**

A) UMAP projection of the dopaminergic dataset with neighborhood color-coding. B) Hierarchical dendrograms with labels and color-coded territories and their respective neighborhoods, as well as ML-clusters. Dotplot shows additional territory (TER) and neighborhood (NH) specific markers.

**Figure 4-figure supplement 1**

**Neighborhoods of *Sox6* territory**

A) Color-coded UMAP projections highlighting the four Sox6 neighborhoods. B-F) Chromogenic in situ hybridization images from Allen Mouse Brain Atlas with the probes indicated, in situ sequencing for *Aldh1a1, Th* and *Ndnf* in D and F. G) Schematic model of the localization of *Sox6* NH1-NH4, following the color code in A), with the most anterior section on the left. Scalebars are 500 µm.

**Figure 4-figure supplement 2**

**Neighborhoods of *Ebf1, Pdia5* and *Otx2* territories**

A-C) Neighborhoods of *Ebf1* territory. *Col24a1* was detected by in situ sequencing, dotted circle indicating area in PBP were *Col24a1*$^{+}$ cells most enriched. *Vip* detected by fluorescent RNA in situ hybridization combined with TH immunohistochemistry. D-F) Neighborhood 2 of *Pdia5* territory showing enriched marker gene expression in the lateral SN. Images in D) from Allen Mouse Brain Atlas. F) Closeup images of SNcL with immunohistochemical staining with antibodies indicated, lateral side towards left. G) Allen Mouse Brain Atlas images for *Otx2* territory markers *Cbln4* and *Lpl*, with the most anterior section upmost. H) Allen Brain Atlas images of *Otx2*_NH1 marker *Grp*, most anterior section upmost. I) In situ sequencing of *Otx2*_NH2 marker *Csf2rb2*, with more anterior section on top. K) Color-coded

UMAP projections of *Ebf1, Pdia5* and *Otx2* territories highlighting the neighborhoods. Scalebars are 200 µm in A, 100 µm in B and F, and 500 µm in D, G-I.

**Figure 4-figure supplement 3**

**Neighborhoods of *Gad2, Fbn2* and *Pcsk6* territories**

A-C) Neighborhoods of *Gad2* territory. A) *Gad2* RNA in situ hybridization in *Pitx3*$^{EGFP/+}$ tissue to detect *Th*$^{low}$ *Gad2* neurons. Arrowheads point to some of the co-expressing cells. C) In situ sequencing with *Ebf2* to detect NH1 and *Met* to detect NH2, with the most anterior section upmost. D-G) Neighborhoods of *Fbn2* territory. D, F) In situ sequencing of *Fbn2, Rxfp1* and *Dsg2*. G) Allen Brain Atlas images of *Rxfp1, Col23a1*, modified by adding lines and text to indicate boundaries of different VTA populations. Arrowheads point to areas where expression was detected. H-I) Neighborhoods of *Pcsk6* territory. Allen Brain Atlas Images of *Pcsk6, Ano2,* and *Cpne2*. In situ sequencing of *Ccdc192*, with arrowhead pointing to areas where expression was seen. B, E, H) Color-coded UMAP projections with neighborhoods highlighted, accompanied by UMAPs showing marker gene expression. Scalebars are 100 µm in A and 500 µm in B, D-G, I.

**Figure 5-figure supplement 1**

**Validation of unilateral 6-OHDA-induced lesion**

A) DAT-binding autoradiography of striatal tissue. See methods for details. B) Quantification of the autoradiographs in A, with both doses of 6-OHDA being significantly different from the intact (un-lesioned) side. C) Mosaic plot showing the residuals of Pearson's Chi square test of independence, with the two conditions as the categorical variables. The plot shows deviations of the observed values from the expected values under the null hypothesis if the categorical variables were independent. However, the test shows that the categorical variables are significantly associated (p-value < 2.2e-16). The negative residuals denote that the observed values are smaller than expected, while the positive residuals denote that the observed values are larger than the expected. The width and height of the mosaics signify the number of nuclei per animal and per condition, respectively. D) Control for non-specific DAT binding with dopamine reuptake inhibitor, nomifensine. E) An example of a 6-OHDA lesioned midbrain, coronal section stained with antibodies against TH and RFP. Scalebar 1 mm.

**Figure 6-figure supplement 1**

**Normalized cell loss per cluster and its correlation with the vulnerability module**

A) Calculated normalized cell loss for dopaminergic-only clusters from the 6 territories shown, visualized as percentages, and color-coded by territory. Section of the hierarchical dendrogram of the same clusters. The vulnerable (black triangle) and resilient (black circle) clusters are marked. See methods for details. Error bars show standard deviation, SD. B) Heatmap of adjusted p values for the pairwise comparisons of cell loss among the same clusters in A. For the Conover-Iman test; p-value = $P(T \geq |t|)$, and null hypothesis ($H_0$) was rejected at $p \leq \alpha/2$, which is 0.025. C, D) Linear regression models, fitted for the normalized cell loss in the specified clusters versus the square root of the vulnerability module scores calculated with and without *Slc6a3*, respectively. The grey bands represent the 95% confidence interval limits. p-value = 3.89e-10 for C, and p-value = 1.73e-09 for D.

**Figure 6-figure supplement 2**

**Vulnerability module scores per cluster for integrated *Th*$^{+}$/*Slc6a3*$^{+}$ dataset**

A) UMAP projection of integrated dataset with territory color-coding, and labels of the clusters. B) Heatmaps of adjusted p values for pairwise comparisons of vulnerability and resilience module scores among the dopaminergic-only clusters, excluding the non-mDA clusters. See methods for the statistical tests used. C-D) Violin plots of vulnerability and resilience module scores for all clusters in the integrated dataset.

**Figure 6-figure supplement 3**

**Co-expression network analysis of lesioned nuclei from mDA neuron territories**

A) Transcriptomically similar nuclei were aggregated into over 4800 Metacells, and their aggregated expression profiles are visualized in two dimensions with UMAP, grouped by their respective territories. Non-mDA clusters are colored gray. B) Dendrogram visualization of the hierarchical clustering of genes into co-expression modules based on the topological overlap matrix (TOM). C) Dotplot of scaled module eigengene (ME) for the 9 calculated modules across mDA territories, with their corresponding abundance per territory. D) The co-expression network of the 9 modules, based on the gene expression of mDA territories from the lesioned side. Genes are shown as nodes, co-expression relationships between genes are shown as edges. The point size is scaled based on the kME values. Intramodular edges are the same color as their corresponding modules, while intermodular edges are colored gray. The strength of the co-expression links between genes is used to scale the opacity of the edges in the network. Network edges are downsampled to 65% for better visualization. The top 20 hub genes and other 5 randomly selected genes per module were used. E) Correlation between each module pair based on their MEs. Modules 2 and 3 are negatively correlated while modules 2 and 8 are positively correlated as highlighted by dashed boxes. F) Boxplot showing average kME values per module. The black line in the box indicates the median while the lower and upper hinges correspond to the first and third quartiles (the 25th and 75th percentiles). The upper / lower whiskers extend from the hinge to the largest / smallest values no further than 1.5 * IQR from the hinge. IQR = the inter-quartile range. G) GO terms plots for the co-expression modules 2 and 3, showing the top 20 terms ranked by p-values. The enrichment analysis was performed on the top 100 genes per module, ranked by kME, using the GO_Biological_Process_2021 database.

**Figure 7-figure supplement 1**

**ML-clusters enriched genes and GO analysis of ML-specific enriched genes**

A) UMAP projection of the dopaminergic dataset with territory color-coding. B) Dotplot with enriched markers for ML clusters, ranked by adjusted p values. C) Venn diagrams with differentially expressed genes in all lesioned nuclei when compared to all intact nuclei, and in ML clusters when compared to all other territories, as well as the common genes between the two sets. D-E) Top 20 enriched GO terms, ranked by p value, using genes upregulated and downregulated only in the ML clusters when compared to all other territories (blue sector in C), with GO_Biological_Process_2021 as the gene-set library.

**Figure 7-figure supplement 2**

**UMAP, Comparative DE analysis and canonical cell type markers in intact-untreated dataset**

A) UMAP projections of the merged intact + untreated nuclei dataset, split by the group, intact vs untreated. B) Venn diagrams showing differentially expressed genes between untreated vs intact nuclei and between lesioned vs intact nuclei as well as the overlap of the two sets of genes. C) UMAP projection of this dataset (intact + untreated), color-coded for the major cell types. D) Markers plotted in UMAP for different cell types in C.

**Figure 8-figure supplement 1**

**Integrated mouse-human mDA dataset**

A, B) Mapped and annotated human dopaminergic control nuclei (B) by using mouse intact nuclei from mDA territories (A) used as reference UMAP. C) Human nuclei in B, split by the transferred labels. D) Numbers of mapped and annotated human nuclei in C, per transferred labels. E, F) Mapped and annotated human dopaminergic control nuclei which were already labeled as Sox6 in C, by using mouse intact nuclei from only Sox6 territory used as reference UMAP. G) Human nuclei in F, split by the transferred labels. H) Numbers of mapped and annotated human nuclei in G, per transferred labels.

**Supplementary Results related to Figure 4-figure supplements 1-3**

***Sox6* territory**

Generally, SNc and its subdomains belonged to the *Sox6* territory, with the exception of SNcL which contained few, if any, *Sox6*$^{+}$ cells, and which belonged to the *Pdia5* territory (see below). *Sox6* is known to be expressed in the adult ventral SNc, but also in some cells of the VTA (Panman et al., 2014). The most ventral *Sox6*$^{+}$ neurons in SNc co-express *Aldh1a1*, whereas the dorsal tier contains *Aldh1a1*$^{-}$ *Sox6*$^{+}$ cells (Panman et al., 2014, Pereira Luppi et al., 2021). Similarly, in our dataset the four neighborhoods of *Sox6* territory could be divided into two groups based on *Aldh1a1* expression (Figure 4B, Supplementary Figure 4): NH1 and NH4, which were entirely *Aldh1a1*$^{+}$, and NH2 and NH3, which were mostly *Aldh1a1*$^{-}$. *Sox6*$^{+}$ *Aldh1a1*$^{+}$ cells have been reported to express *Anxa1*, *Aldh1a7*, and *Lmo3* and project mainly to dorsolateral caudate putamen (Poulin et al., 2014, 2018, 2020, La Manno et al., 2016, Hook et al., 2018, Saunders et al., 2018, Azcorra et al., 2023). Although *Lmo3* was found in all *Sox6* neighborhoods, it was indeed most enriched in both NH1 and NH4 (Supplementary Figure 3).

NH1 consisted mostly of *Anxa1*$^{+}$ neurons, localizing in the SNcM and SNcV (Supplementary Figure 4C, G). *Anxa1* was also found in NH4 which, based on the expression of *Aldh1a7*$^{+}$ and *Grin2c*$^{+}$, most likely matches the most ventral tier of anterior SNc (Supplementary Figure 4D,G). The *Anxa1*$^{+}$ SNpc neurons were recently characterized in a series of behavioral experiments and appear to respond specifically to acceleration (Azcorra et al., 2023). However, as so many genes were shared between NH1 and NH4, the exact location of these neighborhoods in the tissue can only be estimated at this point. More definitive localization will require further analyses.

*Sox6*$^{+}$ *Aldh1a1*$^{-}$ NH2 enriched markers included *Col11a1* and *Vcan,* although *Vcan* was also found in *Aldh1a1*$^{+}$ *Sox6*$^{+}$ cells of NH1, supporting earlier findings (Saunders et al., 2018, Poulin et al., 2020). Allen Brain Atlas expression data for these genes showed localization mostly in the anterior SNc, corresponding to upper, *Aldh1a1*$^{-}$ tier (Supplementary Figure 4E, G). A small population of NH2 cells express also low levels of *Calb1*, which is more highly expressed in VTA (see below). Projections of these *Aldh1a1*$^{-}$ *Calb1*$^{+}$ *Sox6*$^{+}$ neurons in SNpc are biased towards ventromedial regions of caudate putamen (Poulin et al., 2018).

NH3 was mostly *Aldh1a1*$^{-}$ *Ndnf*$^{+}$ *Vill*$^{+}$, although both *Vill* and *Ndnf* were also found to some extent in NH1 and NH4 (Supplementary Figure 4F, G). Our ISS sequencing and Allen Brain Atlas data showed that *Ndnf*$^{+}$ *Aldh1a1*$^{-}$ neurons in more dorsal tier of SNc extended through the midbrain, and *Vill* in Allen Brain Atlas was detected in the same areas, although in fewer cells. NH3 expressed also *Igf1*, shown to be expressed in both anterior and caudal SNc (Supplementary Figure 3, Pristera et al., 2018). Furthermore, this neighborhood expressed *Tacr3* (Figure 3C, Supplementary Figure 3), which together with *Ndnf* had been reported to be expressed in *Sox6*$^{+}$ *Aldh1a1*$^{-}$ neurons which project to striatum (Poulin et al.,

2014, 2018, 2020), although neurons with similar expression profile were found also in *Pdia5* territory (see below).

One cluster in NH3 did express *Aldh1a1* (Figure 3C), which likely represents *Aldh1a1*$^{+}$ cells detected in more caudal SNc, where neither NH1 nor NH4 -specific markers, such as *Aldh1a7* or *Anxa1*, were found (Supplementary Figure 4B, D). Some of these neurons likely correspond to *Sox6*$^{+}$ neurons projecting to the core and lateral shell of nucleus accumbens (Poulin et al., 2018).

##### *Ebf1* territory

*Ebf1*, the territory with fewest cells, consisted of two neighborhoods (Figure 4A, F, Supplementary Figure 5A-C). NH1, specified by *Col24a1*, was found in scattered cells of PAG, RLi, and PBP, although here the ISS signal was diffuse and hard to interpret (Supplementary Figure 5A). This neighborhood, which expressed also *Slc17a6, Calb1, Cck, Npw* and *Man1a*, likely corresponds to a specific population of *Npw*$^{+}$ DA neurons identified in PAG, VTA and dorsal raphe, which project to central extended amygdala and modulate mouse behavior under stress (Motoike et al., 2016).

NH2 expressed *Vip* and these cells were found mostly in periaqueductal grey (PAG) and DR, and to some extent in PBP, PN, RLi and CLi (Supplementary Figure 5B). This neighborhood corresponds to the previously identified *Slc17a6*$^{+}$ *Vip*$^{+}$ subtype (Dougalis et al., 2012, Poulin et al., 2014, La Manno et al., 2016, Hook et al., 2018, Tiklova et al., 2019). Vip-Cre- labelled neurons project to lateral part of the central amygdala (Poulin et al., 2018).

##### *Pdia5* territory

Many of the cells in *Pdia5* territory were also *Sox6*$^{+}$ (Figure 3C), and may correspond to the *Sox6*$^{+}$ *Aldh1a1*$^{-}$ subtype that has been detected in PBP and in the most dorsal tier of SNc

(Poulin et al., 2014, La Manno et al., 2016) expressing *Igf1*$^+$, *Ndnf*$^+$ and *Tacr3*$^+$. Based on these markers, it matches *Pdia5* NH1 (Figure 3C, Supplementary Figure 3). *Pdia5* itself showed an interesting expression pattern in VTA, forming a distinct dorsal tier of PBP which extended laterally as a narrow strip of cells above SNc, divided by medial lemniscus (Figure 4C, F). A careful chemoarchitectonic and cell morphological analyses of midbrain DA neurons by Paxinos and colleagues assign this dorsalmost tier as a part of PBP, not as a part of SNc proper (Fu et al., 2011). Furthermore, the *Sox6*$^+$ neurons in this layer project to ventromedial parts of striatum, as opposed to the rest of SNc which mainly projects to dorsolateral striatum (Pereira Luppi et al, 2021).

Although *Pdia5* NH1 and NH2 shared many genes, the expression of *Postn* and *Npy1r* in NH2 enabled us to map the majority of cells in this neighborhood to the most lateral tip of SNcL (Supplementary Figure 5D, E), which was also *Calb1*$^+$ (Supplementary Figure 5F). In addition, this neighborhood expressed *Slc17a6, Cbln1,* and *Wnt7b* (Figure 3C, Supplementary Figure 3) and likely represents some of *Slc17a6*$^+$ mDA neurons described earlier (Poulin et al., 2020). These neurons innervate mainly the central nucleus of amygdala and the tail of caudate putamen (Poulin et al., 2018).

**Otx2 territory**

*Otx2*$^+$ *Aldh1a1*$^+$ DA neurons in VTA have been described before (Di Salvio et al., 2010). In addition to *Otx2*, the previously identified enriched genes for this subtype include *Lpl, Tacr3, Grp,* and *Cbln4* (Poulin et al., 2014, La Manno et al., 2016, Tiklova et al., 2018, Hook et al., 2018, Saunders et al., 2018). We found these genes to be enriched in the *Otx2* territory, and based on our own analyses and Allen Mouse Brain Atlas data, the cells of this territory appears to be localized in the most medial regions of VTA: IF, PN, medioventral PBP, as well is to some extent in RLi and CLi (Figure 4D, F, Supplementary Figure 5G), which supported the earlier analyses.

*Otx2* NH1 was *Tacr3*$^{+}$ *Aldh1a1*$^{+}$ *Slc6a3*$^{high}$ and also expressed *Grp* and *NeuroD6*, which specify a distinct VTA subtype (Kramer et al., 2018). *Grp*$^{+}$ *NeuroD6*$^{+}$ cells innervate the medial shell of the nucleus accumbens, whereas Grp$^{+}$ NeuroD6$^{-}$ cells send projections to the dorsomedial striatum (Kramer et al., 2018). Aldh1a1-Cre labelled cells in the VTA, corresponding to *Otx2* NH1, have been shown to innervate the medial shell of nucleus accumbens and olfactory tubercle (Poulin et al., 2018). NH1 cells were found in PN, PIF and the most medial parts of PBP (Supplementary Figure 5H). The *Aldh1a1*$^{-}$ *Slc6a3*$^{low}$ NH2 cells, which also expressed *Csf2rb2,* were mostly localized in RLi and IF (Supplementary Figure 5I). *Slc6a7 (Vglut2)* was expressed in both NH1 and NH2, although being more highly expressed in NH2. Vglut2-Cre-labelled *Otx2*$^{+}$ cells in ventromedial VTA innervate mainly nucleus accumbens medial shell and in discrete areas of olfactory tubercle (Poulin et al., 2018). It should be noted that *Slc6a7* is also found in *Ebf1* and *Gad2* territories as well as in *Pdia5* NH2. However, this neighborhood also expresses *Cck* – which is also found in *Ebf1* territory, *Pcsk6* NH2, and *Pdia5* NH1 – and Cck-Cre-labelled neurons in VTA project to basolateral amygdala (Poulin et al., 2018).

***Gad2* territory**

*Slc32a1*$^{+}$ *Slc17a6*$^{+}$ *Slc6a3*$^{low}$ *Th*$^{low}$ DA neurons in VTA have been identified in several studies (Poulin et al., 2014, La Manno et al., 2016, Tiklova et al., 2018, Saunders et al., 2018), and based on the enriched gene expression match our *Gad2* territory. In support of these earlier studies, we localized these neurons mostly to IF to some extent RLi, CLi and PBP (Figure 4A, F, Supplementary Figure 6A-C). Both *Ebf2*$^{+}$ NH1 and *Met*$^{+}$ NH2 were intermingled in these areas, although NH2 cells concentrated more in the ventral midline of

RLi and were less frequent in more caudal VTA (Supplementary Figure 6C). These neurons likely correspond to previously described VTA neurons that project to the lateral habenula and function to promote award (Stamatakis et al., 2013). Although the majority of neurons in this territory express low levels of both *Th* and *Slc6a3*, they do express the typical mDA neuron signature, suggesting a potential for GABA and DA co-release.

##### *Fbn2* territory

The *Fbn2* territory consisted of two small neighborhoods, both of which were localized in the antero-medial PBP (Figure 4A, F, Supplementary Figure 6D-G). In addition to the most specific territory marker *Fbn2*, these cells expressed other less specific identifiers such as *Slc17a6, Mkx, Mid1, Glra3* and *Otx2* (Figure 3, Supplementary Figure 3) and some of them likely corresponded to *Slc17a6*$^{+}$ mDA subtype in VTA (Poulin et al., 2020, Garritsen et al., 2023).

NH1 was further specified by *Rxfp1*, whereas NH2 expressed *Dsg2* and *Col23a1* (Supplementary Figure 6F, G). Another highly enriched, although not exclusive, expressed gene in *Fbn2* territory was *Bcl11a* (Supplementary Figure 3), suggesting that some of the recently identified *Bcl11a*$^{+}$ DA neurons in VTA (Tolve et al., 2021) likely belong to this territory. Some of the neurons in *Fbn2*-territory likely match to Vglut2-labelled cells in PBP projecting to nucleus accumbens (Poulin et al., 2018).

##### *Pcsk6* territory

*Pcsk6* territory expressed *Ano2* but had very low *Slc17a6*. *Pcsk6*_NH1 expressed *Ranbp3l, Cgnl1, Cpne2, Ccdc192 (1700011I03Rik)* and *Kank1* (Supplementary Figure 3, Supplementary Figure 6H), and did not correspond well to any mDA neuron types described previously. Our ISS analysis discovered some of *Ccdc192*$^{+}$ cells in PBP where a less exclusive NH1 marker *Cpne2* was also found (Supplementary Figure 6I).

*Pcsk6*_NH2 was best identified by the expression of *Tacr3* and the lack of *Aldh1a1* (Figure 4E). Some of them, which also express *Sox6* and *Ndnf*, match previously described *Sox6*$^{+}$ *Aldh1a1*$^{-}$ *Ndnf*$^{+}$ DA 1B -neurons (Poulin et al., 2014), *Cyp26b1*$^{+}$ *Sox6*$^{+}$ *Tacr3*$^{+}$ mDA subtype 4-3 neurons (Saunders et al., 2018, Poulin et al., 2020) and *Cd9*$^{+}$ *Ntf3*$^{+}$ DA-VTA1 neurons (La Manno et al., 2016). These cells were found in PBP, IF, PN, CLi, and RRF (Figure 4F). Together with *Pdia5* NH1, they formed a dorsolateral extension of PBP.

La Manno G, Gyllborg D, Codeluppi S, Nishimura K, Salto C, Zeisel A, Borm LE, Stott SRW, Toledo EM, Villaescusa JC, Lönnerberg P, Ryge J, Barker RA, Arenas E, Linnarsson S.

2016. Molecular Diversity of Midbrain Development in Mouse, Human, and Stem Cells. *Cell* **167**. doi:10.1016/j.cell.2016.09.027

Motoike T, Long JM, Tanaka H, Sinton CM, Skach A, Williams SC, Hammer RE, Sakurai T, Yanagisawa M. 2016. Mesolimbic neuropeptide W coordinates stress responses under novel environments. *Proc Natl Acad Sci U S A* **113**. doi:10.1073/pnas.1518658113

Panman L, Papathanou M, Laguna A, Oosterveen T, Volakakis N, Acampora D, Kurtsdotter I, Yoshitake T, Kehr J, Joodmardi E, Muhr J, Simeone A, Ericson J, Perlmann T. 2014. Sox6 and Otx2 control the specification of substantia nigra and ventral tegmental area dopamine neurons. *Cell Rep* **8**. doi:10.1016/j.celrep.2014.07.016

Pereira Luppi M, Azcorra M, Caronia-Brown G, Poulin JF, Gaertner Z, Gatica S, Moreno-Ramos OA, Nouri N, Dubois M, Ma YC, Ramakrishnan C, Fenno L, Kim YS, Deisseroth K, Cicchetti F, Dombeck DA, Awatramani R. 2021. Sox6 expression distinguishes dorsally and ventrally biased dopamine neurons in the substantia nigra with distinctive properties and embryonic origins. *Cell Rep* **37**. doi:10.1016/j.celrep.2021.109975

Poulin, J. F., Caronia, G., Hofer, C., Cui, Q., Helm, B., Ramakrishnan, C., Chan, C. S., Dombeck, D. A., Deisseroth, K., & Awatramani, R. (2018). Mapping projections of molecularly defined dopamine neuron subtypes using intersectional genetic approaches. *Nature Neuroscience*, *21*(9). https://doi.org/10.1038/s41593-018-0203-4

Poulin JF, Gaertner Z, Moreno-Ramos OA, Awatramani R. 2020. Classification of Midbrain Dopamine Neurons Using Single-Cell Gene Expression Profiling Approaches. *Trends Neurosci*. doi:10.1016/j.tins.2020.01.004

Poulin JF, Zou J, Drouin-Ouellet J, Kim KYA, Cicchetti F, Awatramani RB. 2014. Defining midbrain dopaminergic neuron diversity by single-cell gene expression profiling. *Cell Rep* **9**. doi:10.1016/j.celrep.2014.10.008

Saunders A, Macosko EZ, Wysoker A, Goldman M, Krienen FM, de Rivera H, Bien E, Baum M, Bortolin L, Wang S, Goeva A, Nemesh J, Kamitaki N, Brumbaugh S, Kulp D, McCarroll SA. 2018. Molecular Diversity and Specializations among the Cells of the Adult Mouse Brain. *Cell* **174**. doi:10.1016/j.cell.2018.07.028

Stamatakis AM, Jennings JH, Ung RL, Blair GA, Weinberg RJ, Neve RL, Boyce F, Mattis J, Ramakrishnan C, Deisseroth K, Stuber GD. 2013. A Unique Population of Ventral Tegmental Area Neurons Inhibits the Lateral Habenula to Promote Reward. *Neuron* **80**:1039–1053. doi:10.1016/j.neuron.2013.08.023

Tiklová K, Björklund ÅK, Lahti L, Fiorenzano A, Nolbrant S, Gillberg L, Volakakis N, Yokota C, Hilscher MM, Hauling T, Holmström F, Joodmardi E, Nilsson M, Parmar M, Perlmann T. 2019. Single-cell RNA sequencing reveals midbrain dopamine neuron diversity emerging during mouse brain development. *Nat Commun* **10**. doi:10.1038/s41467-019-08453-1

Tolve M, Ulusoy A, Patikas N, Islam KUS, Bodea GO, Öztürk E, Broske B, Mentani A, Wagener A, van Loo KMJ, Britsch S, Liu P, Khaled WT, Metzakopian E, Baader SL, Di Monte DA, Blaess S. 2021. The transcription factor BCL11A defines distinct subsets of midbrain dopaminergic neurons. *Cell Rep* **36**. doi:10.1016/j.celrep.2021.109697

### Figure Supplement 1-1

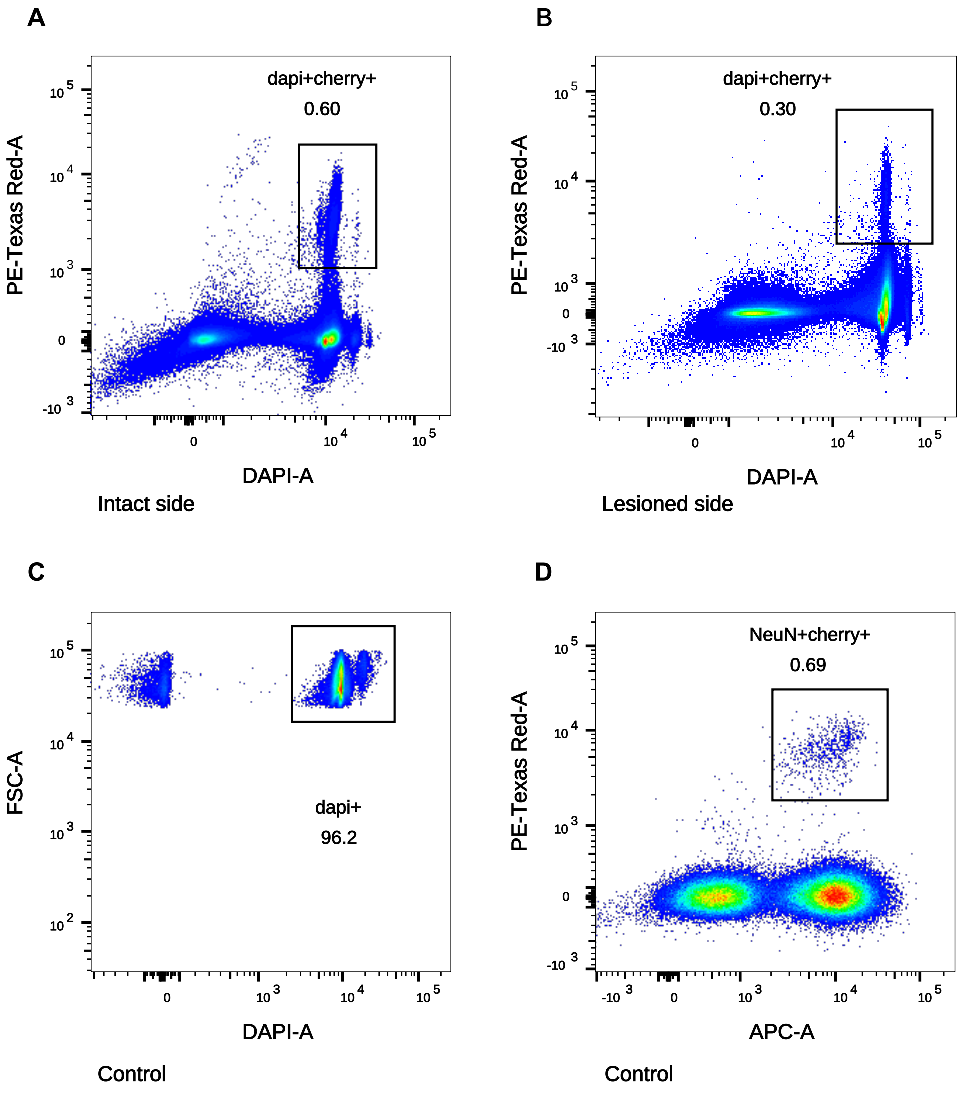

Figure Supplement 1-2

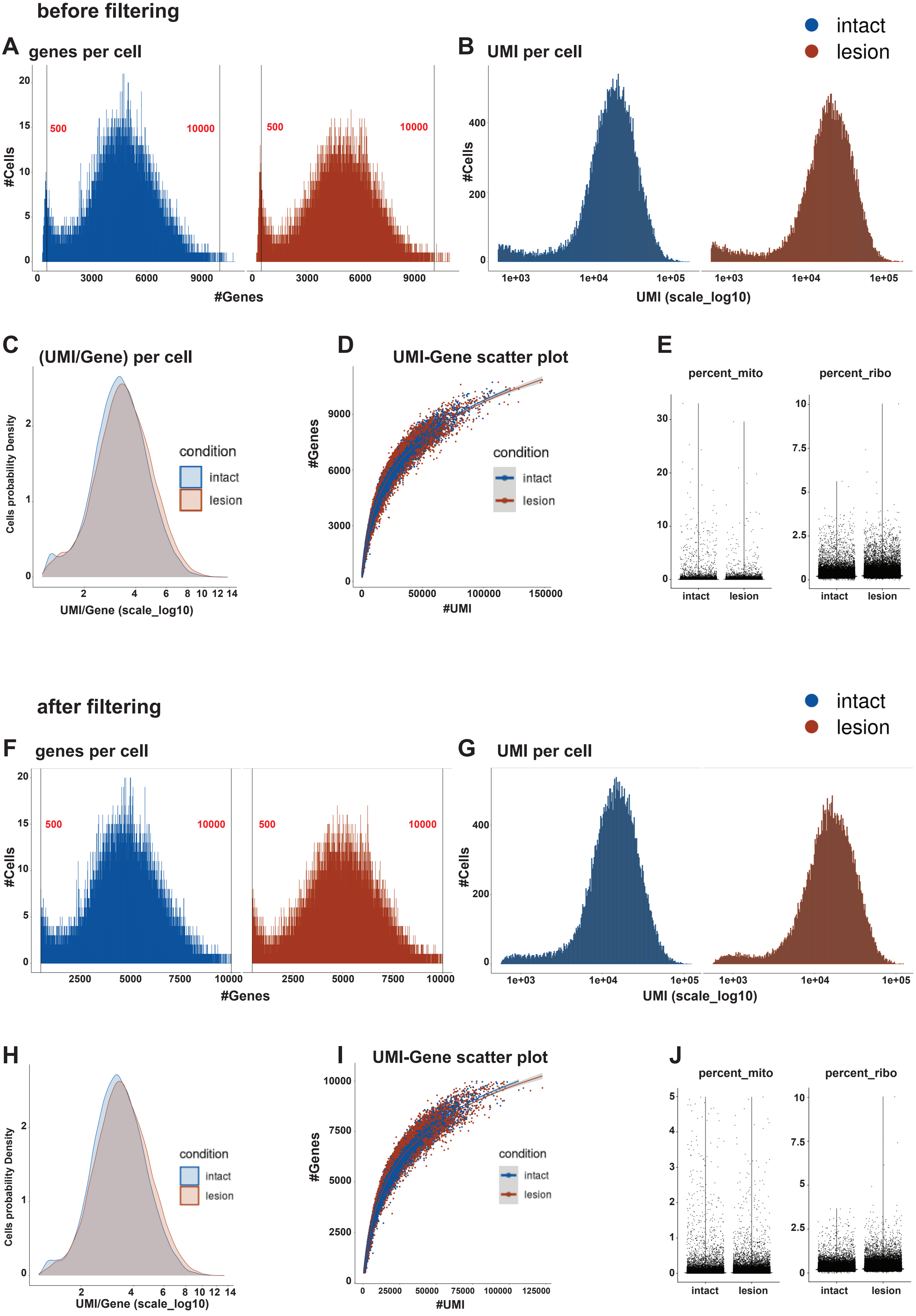

**Figure Supplement 1-3**

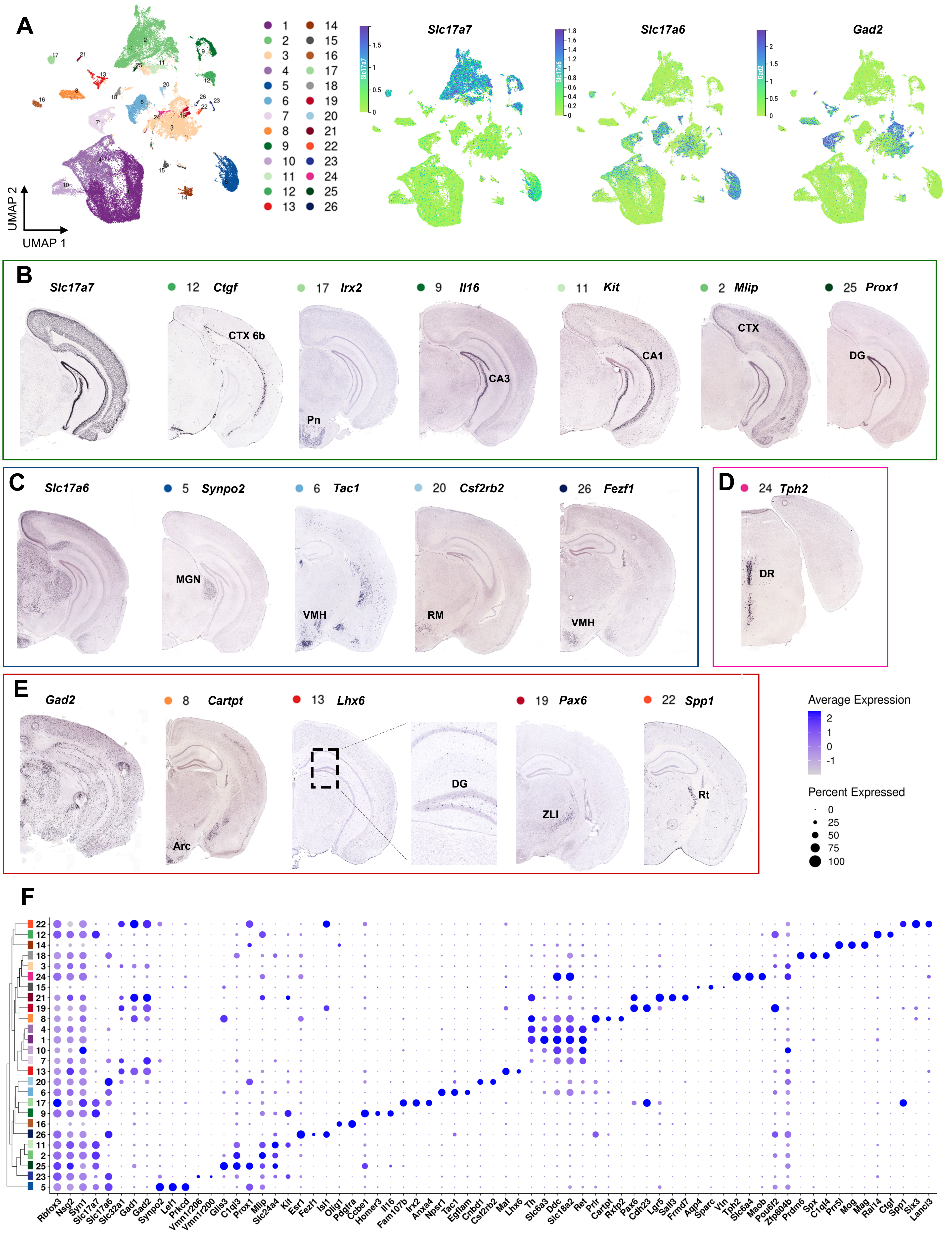

Figure Supplement 2-1

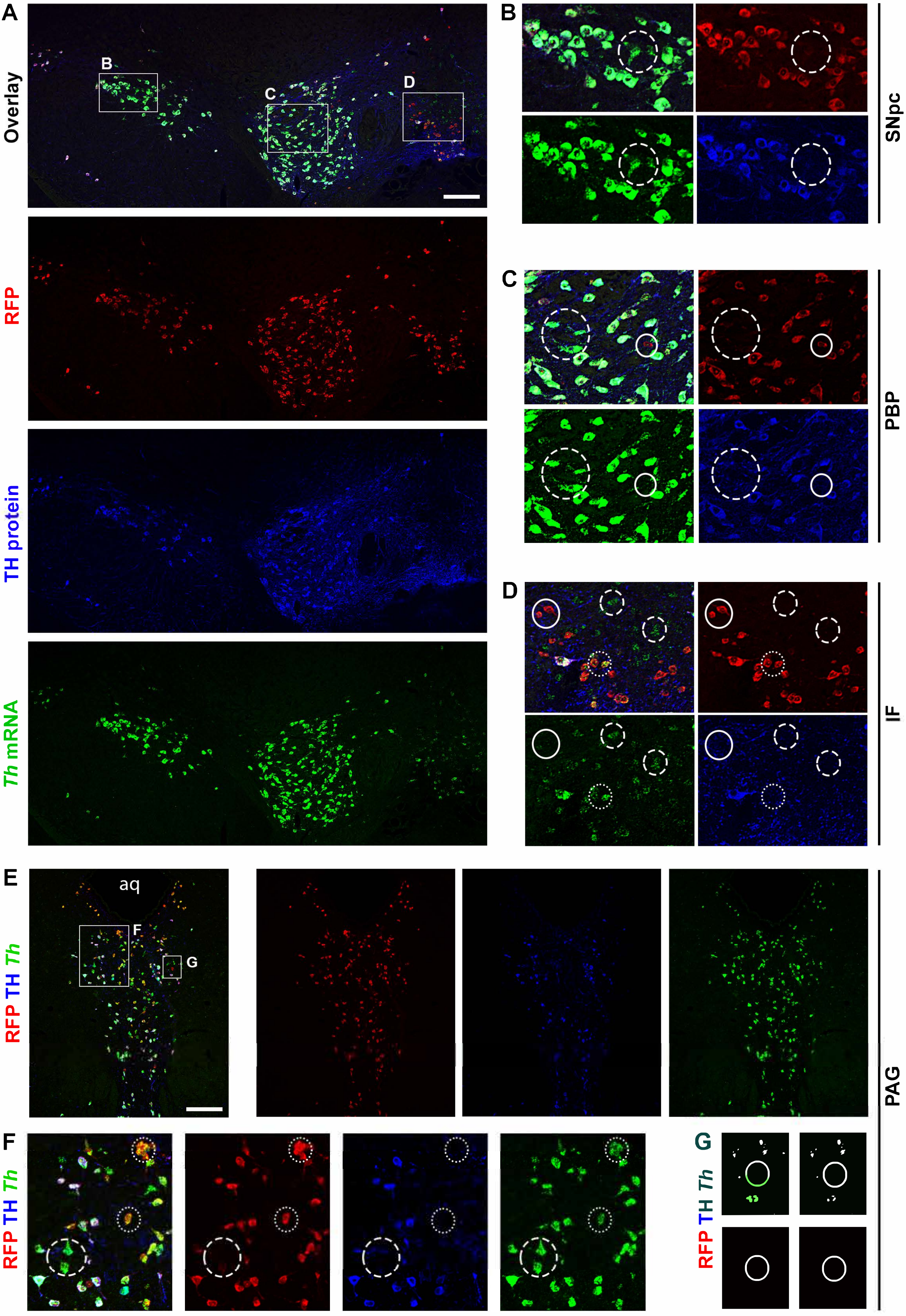


Dat-Cre+ Th mRNA+ TH protein- Dat-Cre- Th mRNA+ TH protein- Dat-Cre+ Th mRNA- TH protein-

#### Figure Supplement 3-1

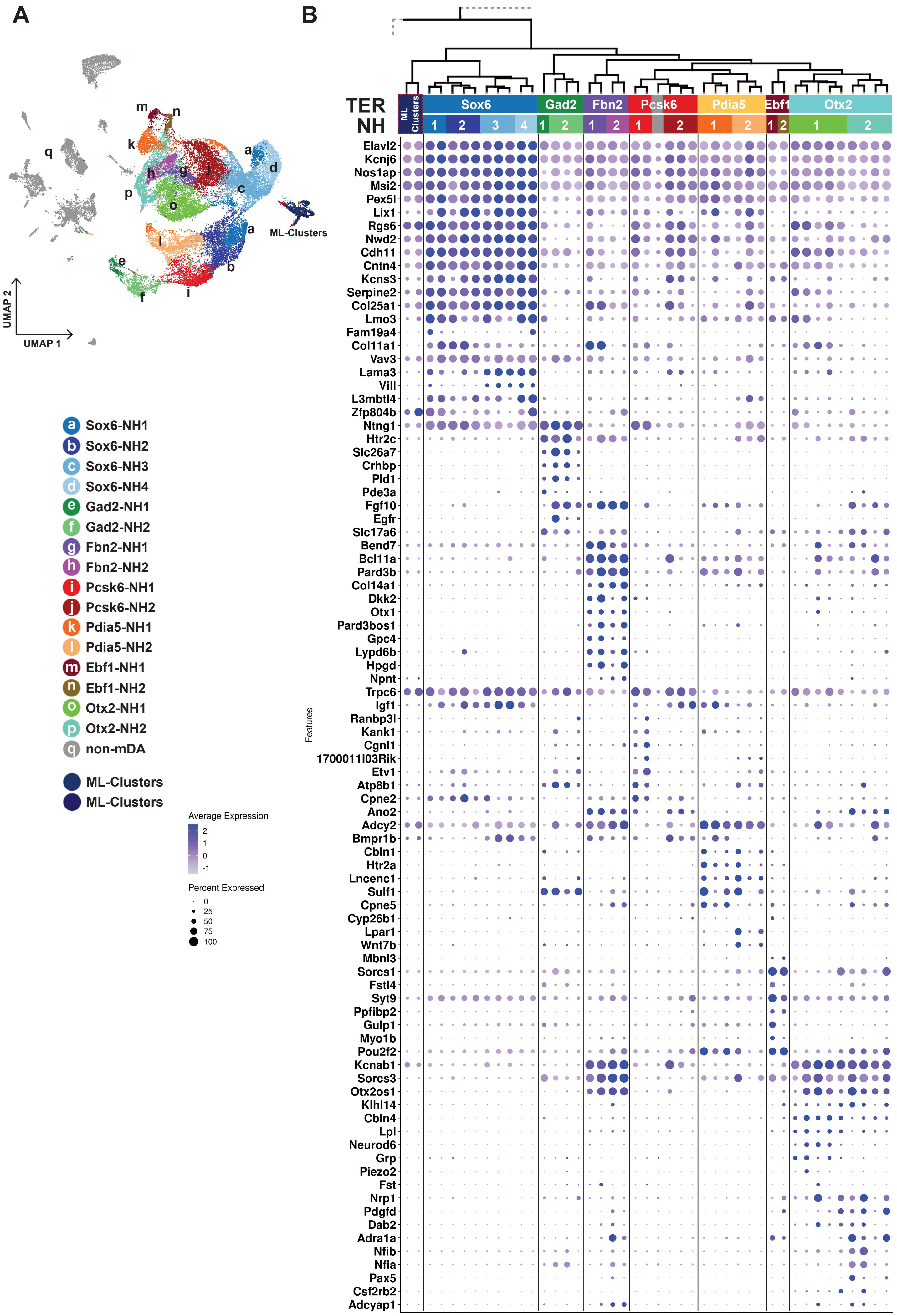

Figure Supplement 4-1

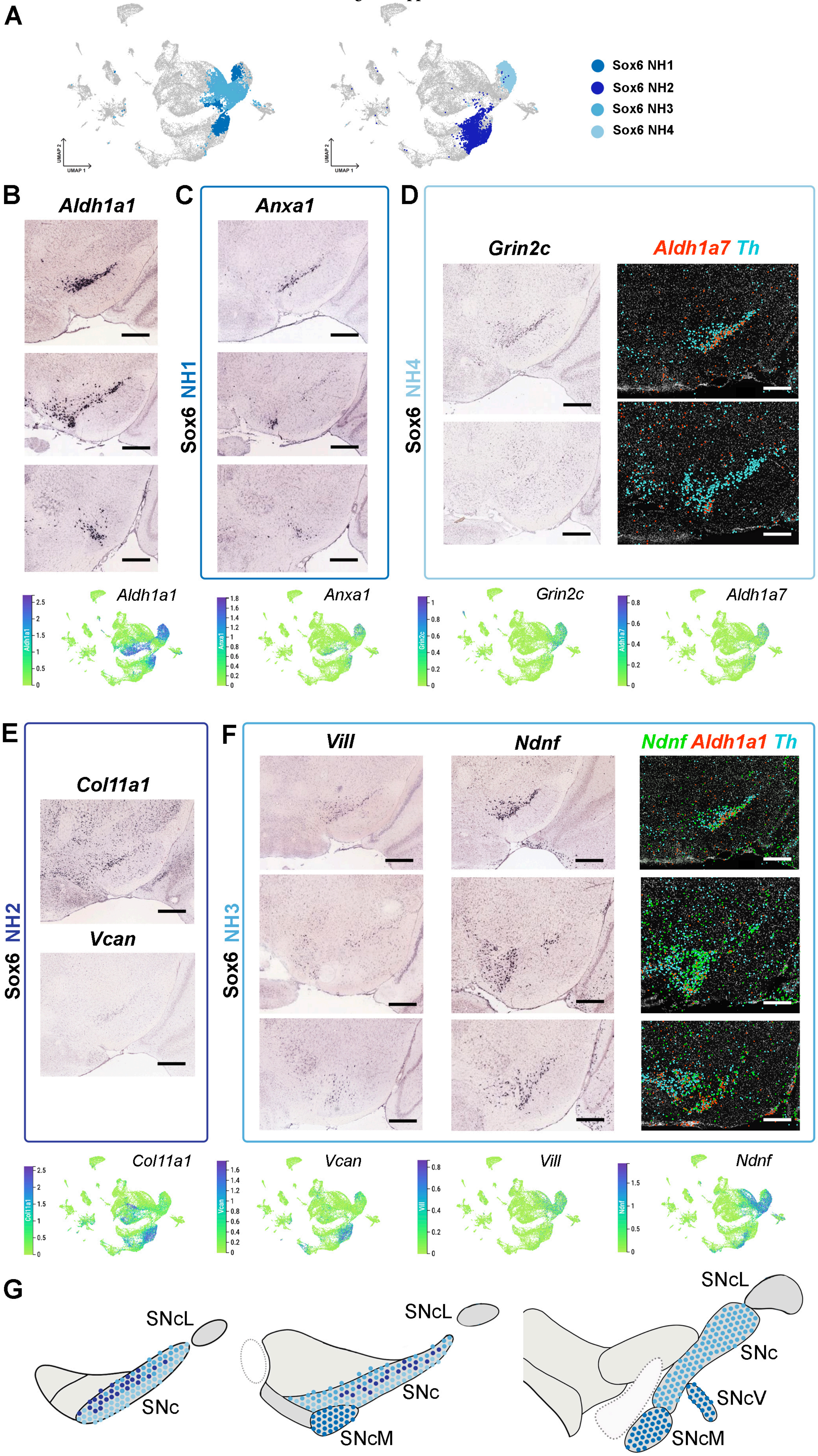

Figure Supplement 4-2

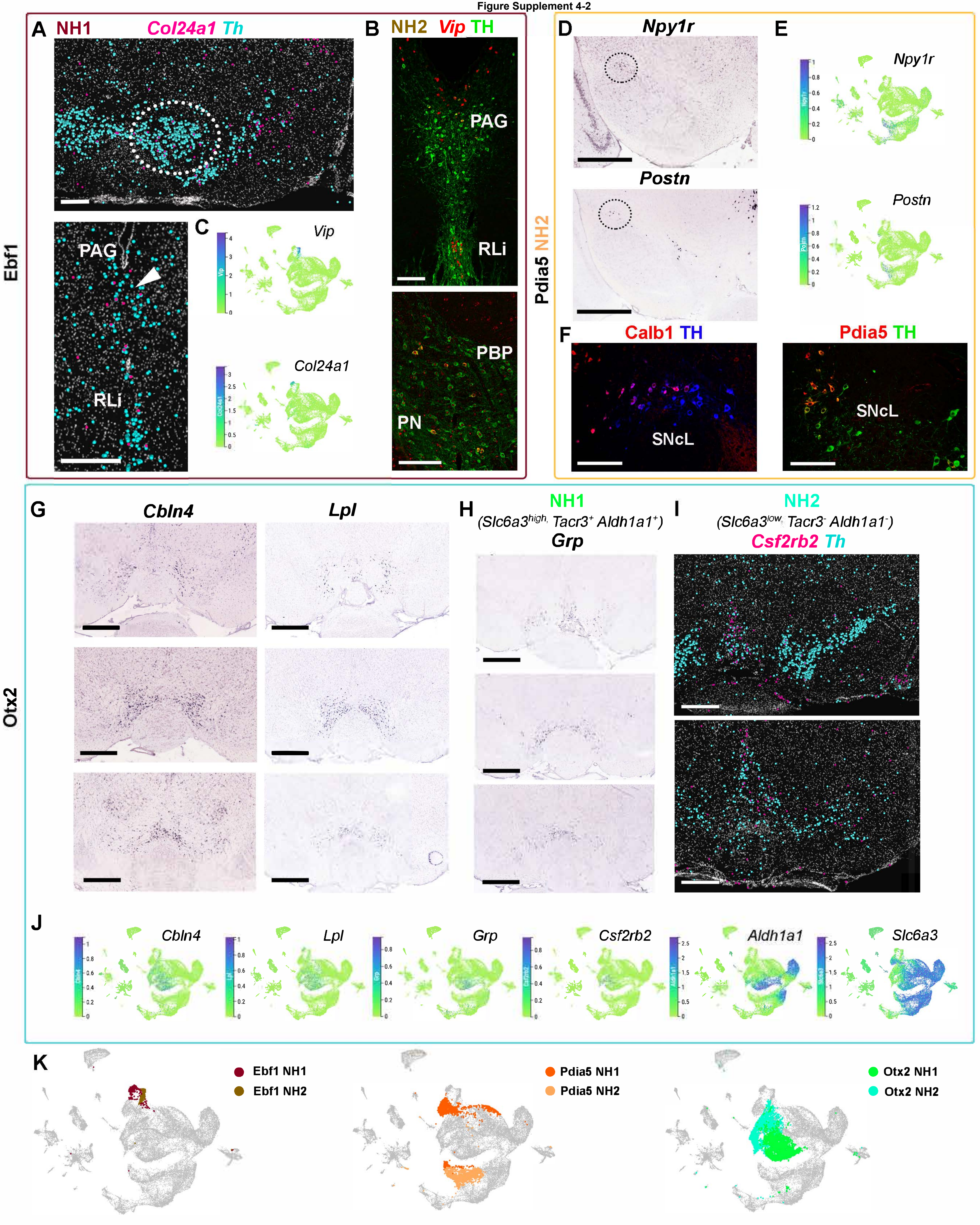

Figure Supplement 4-3

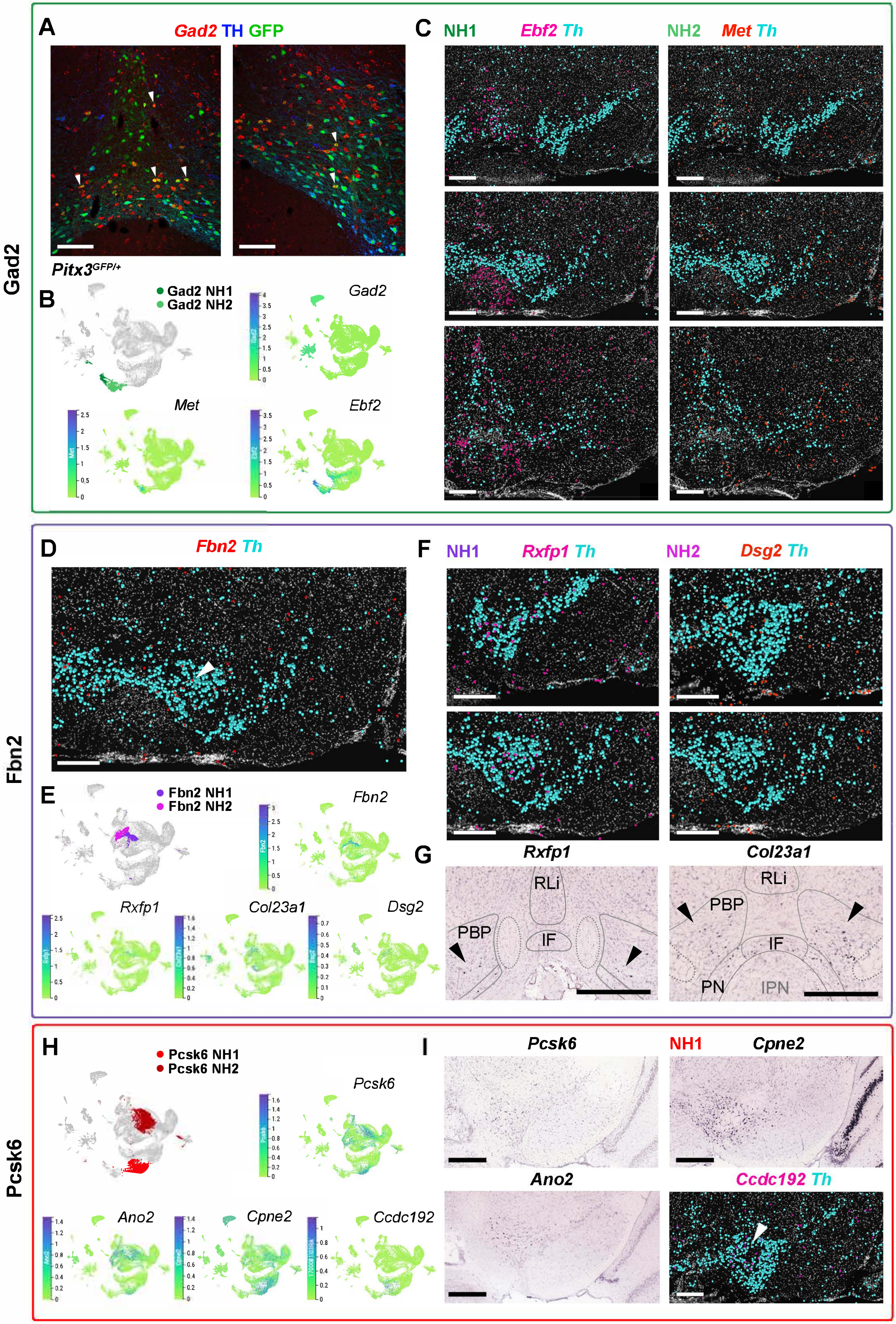

Figure Supplement 5-1

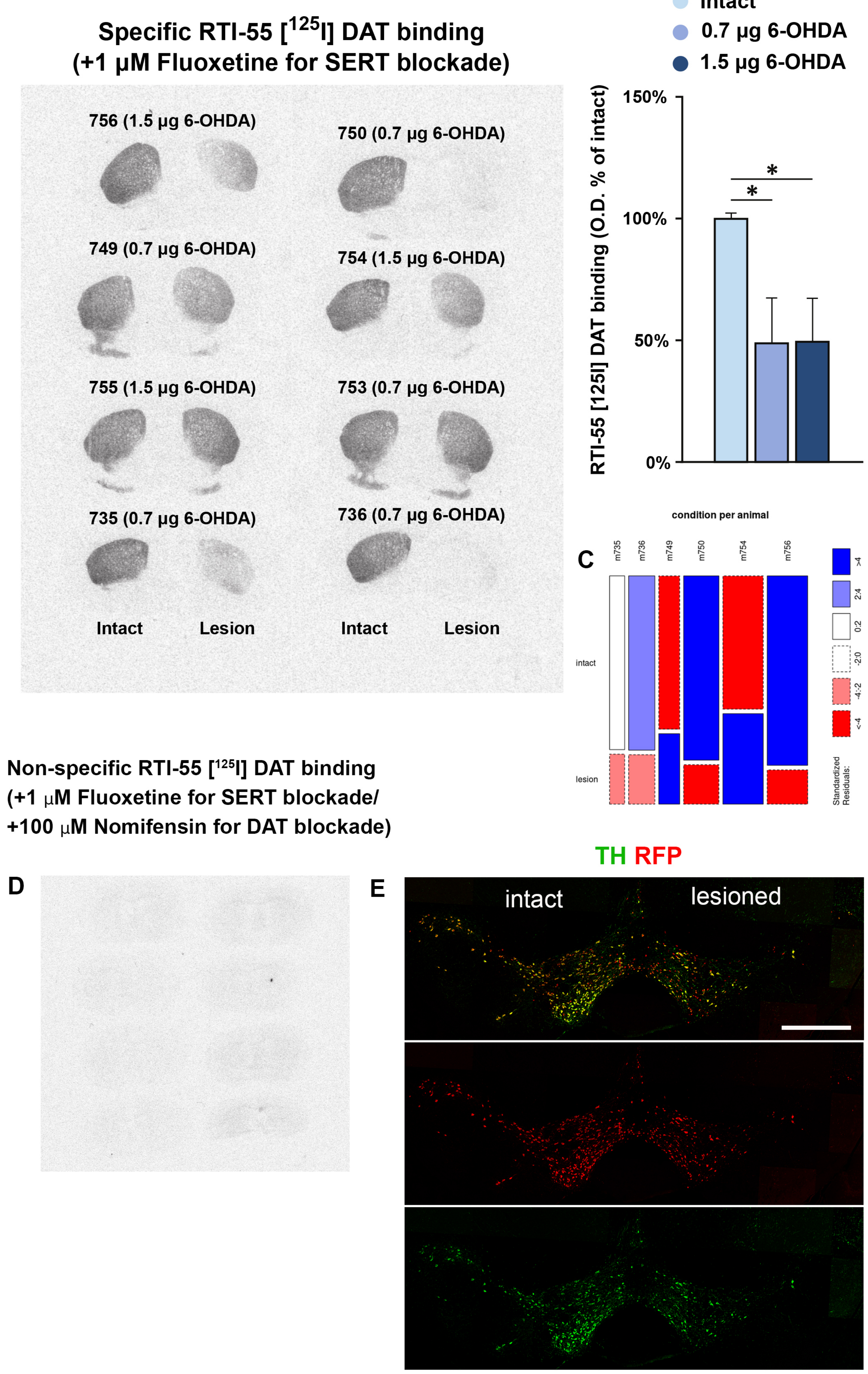

#### Figure Supplement 6-1

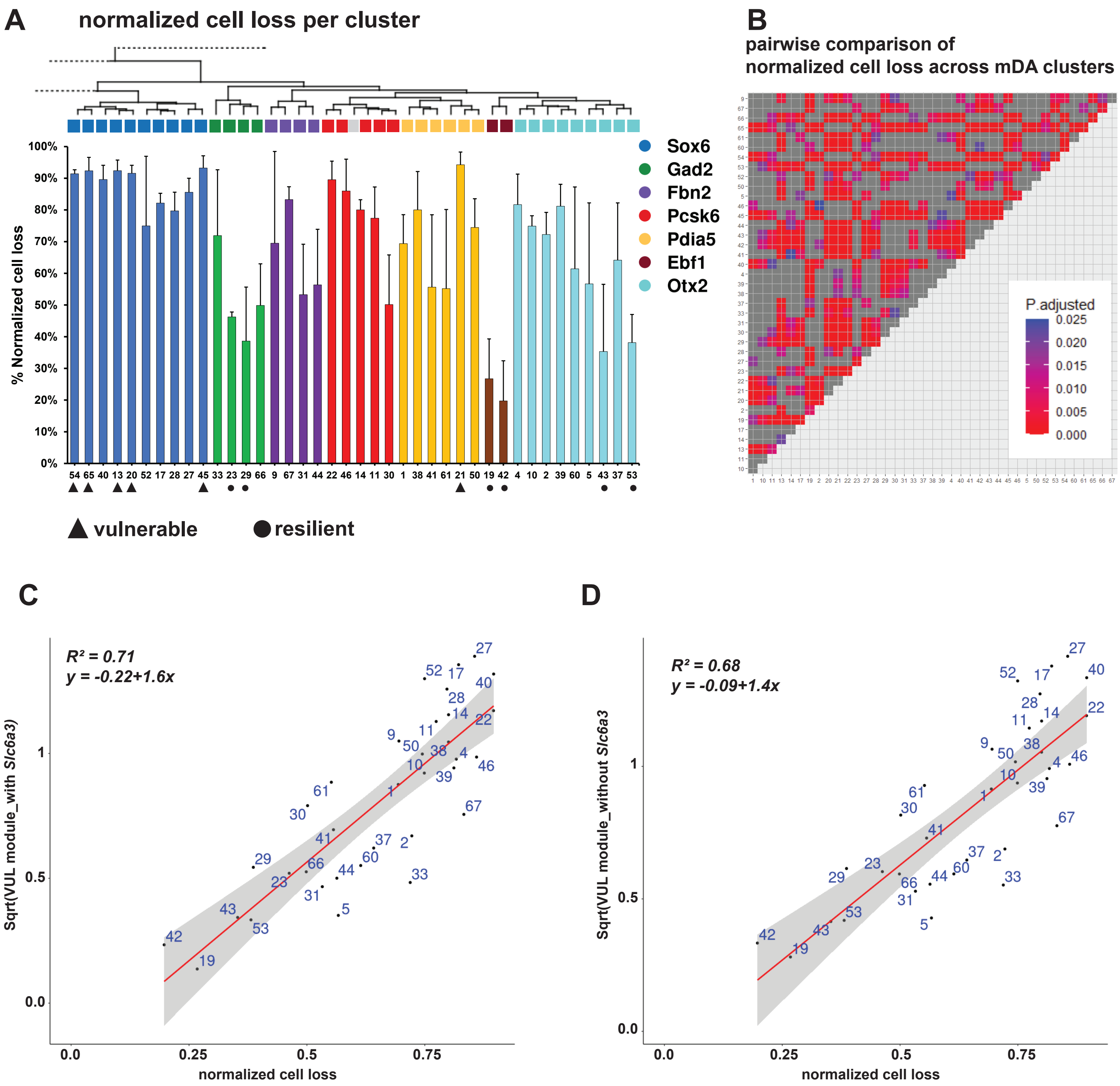

**Figure Supplement 6-2**

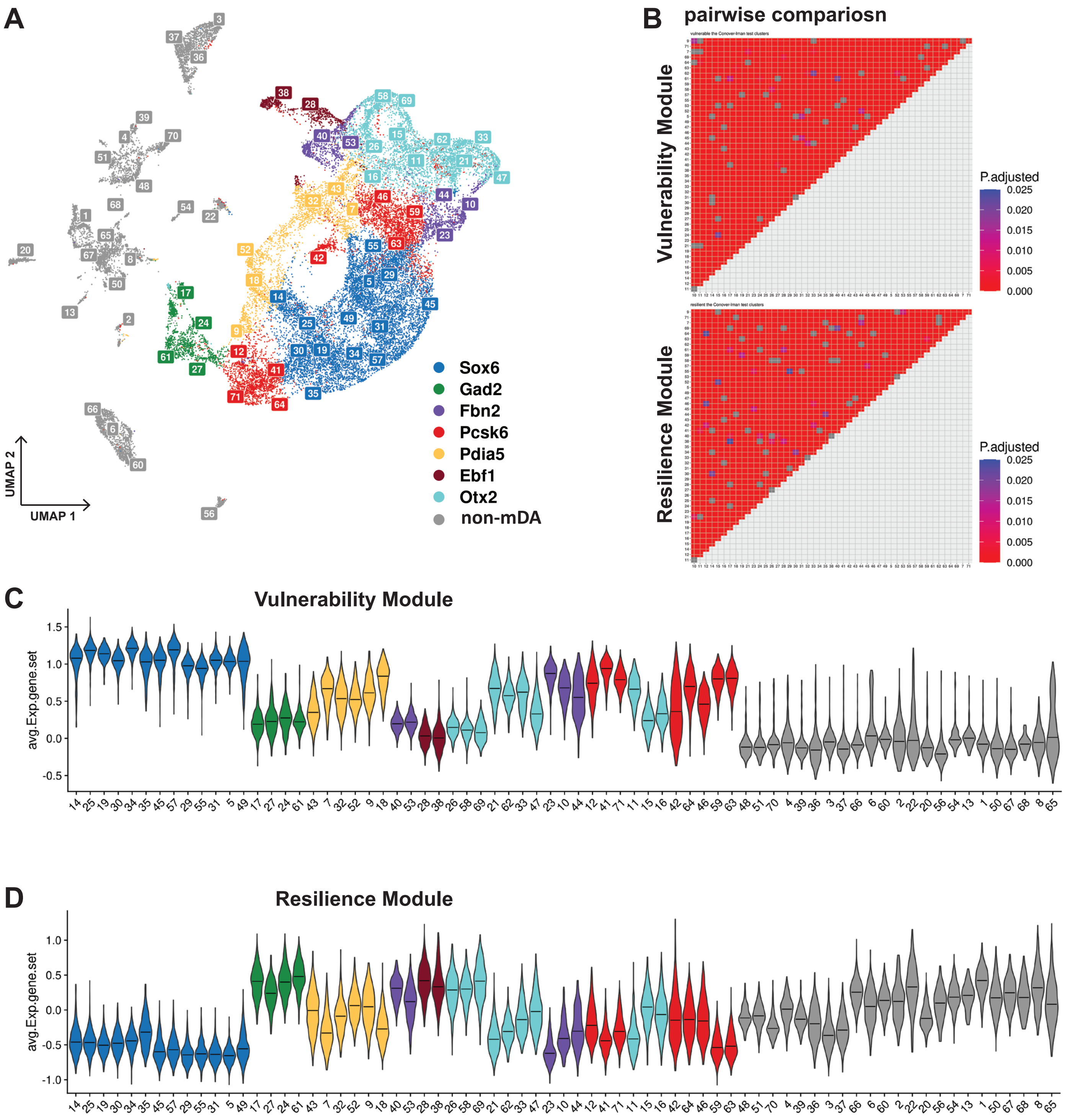

### Figure Supplement 6-3

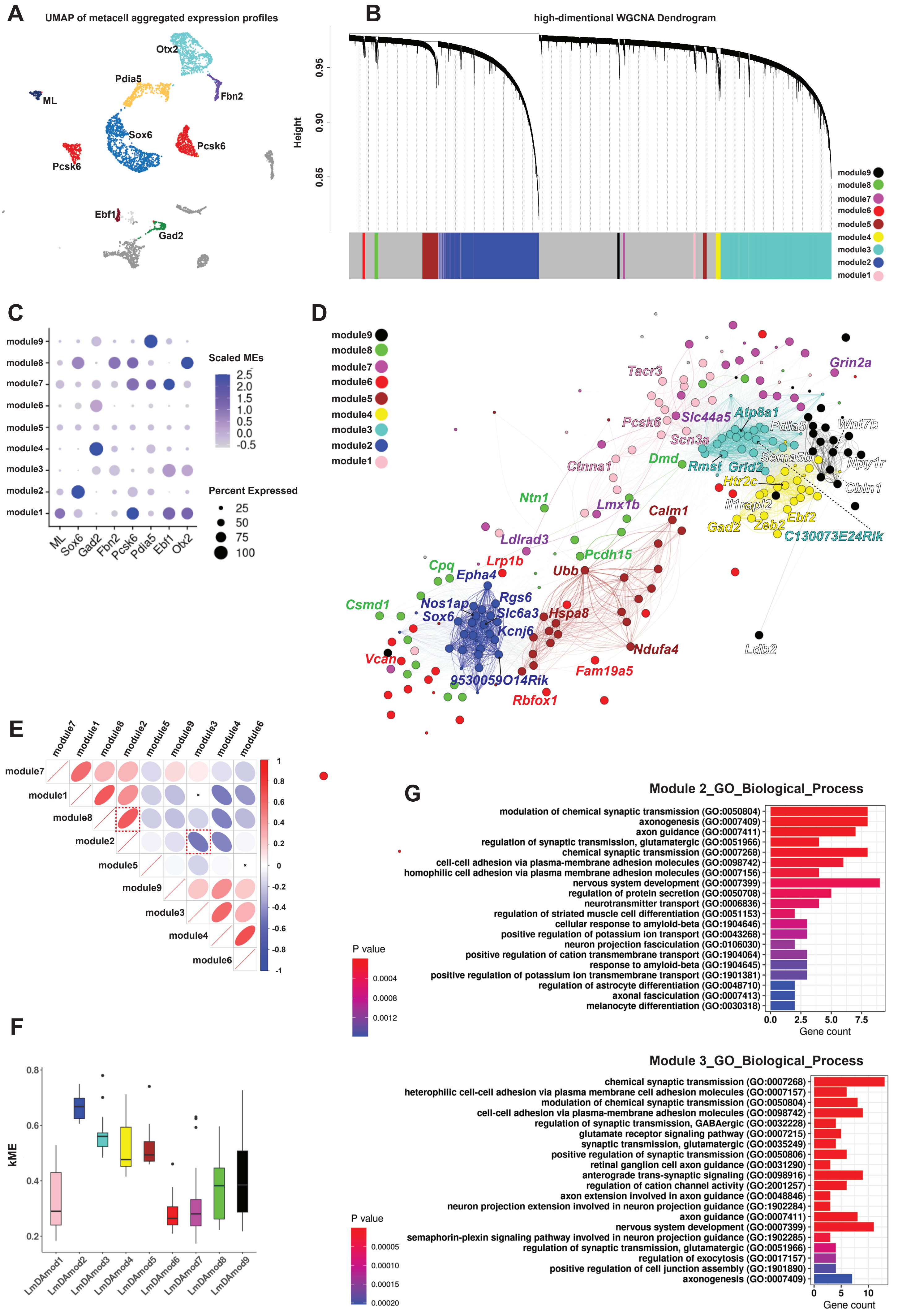

### Figure Supplement 7-1

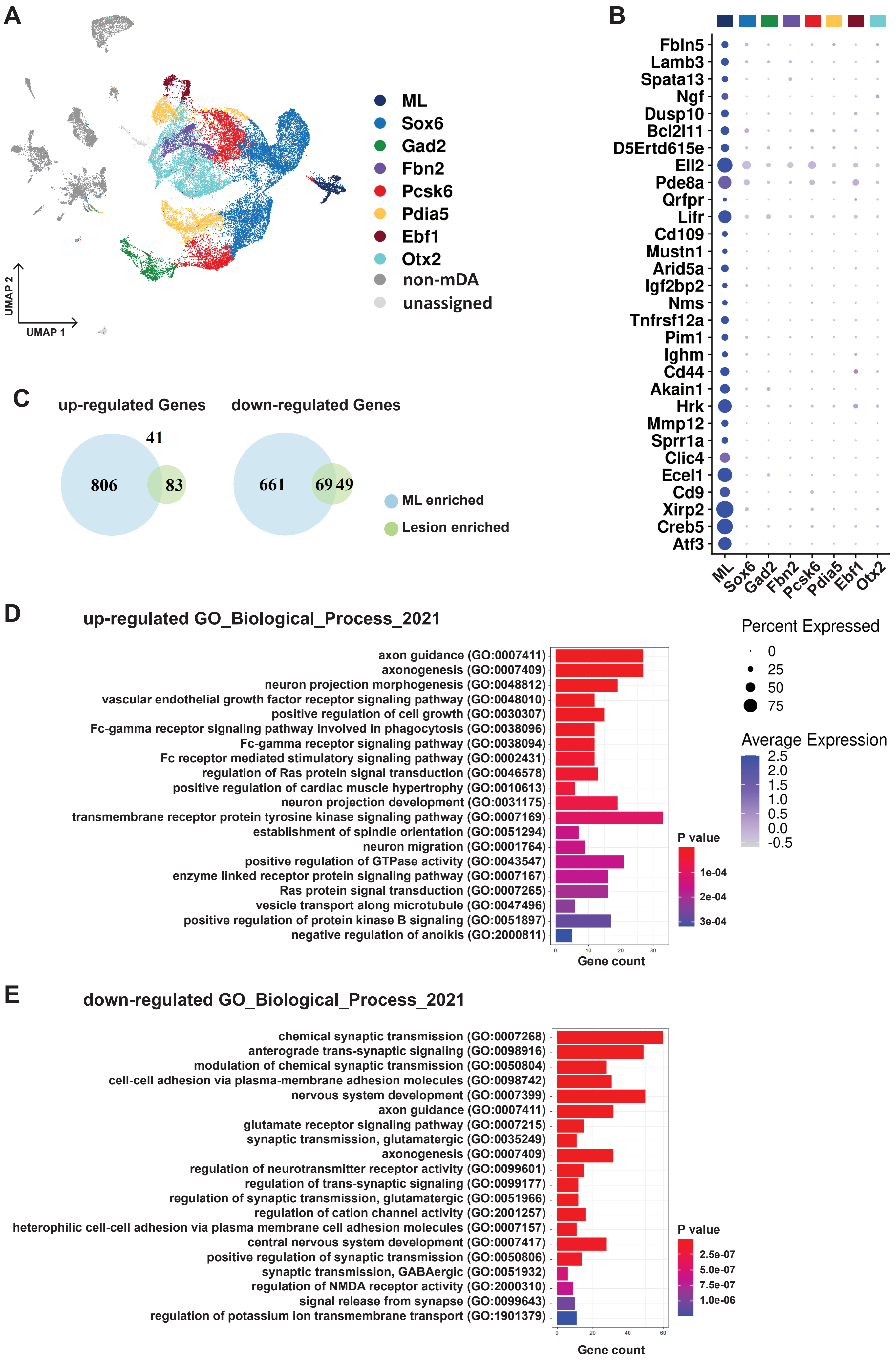

#### Figure Supplement 7-2

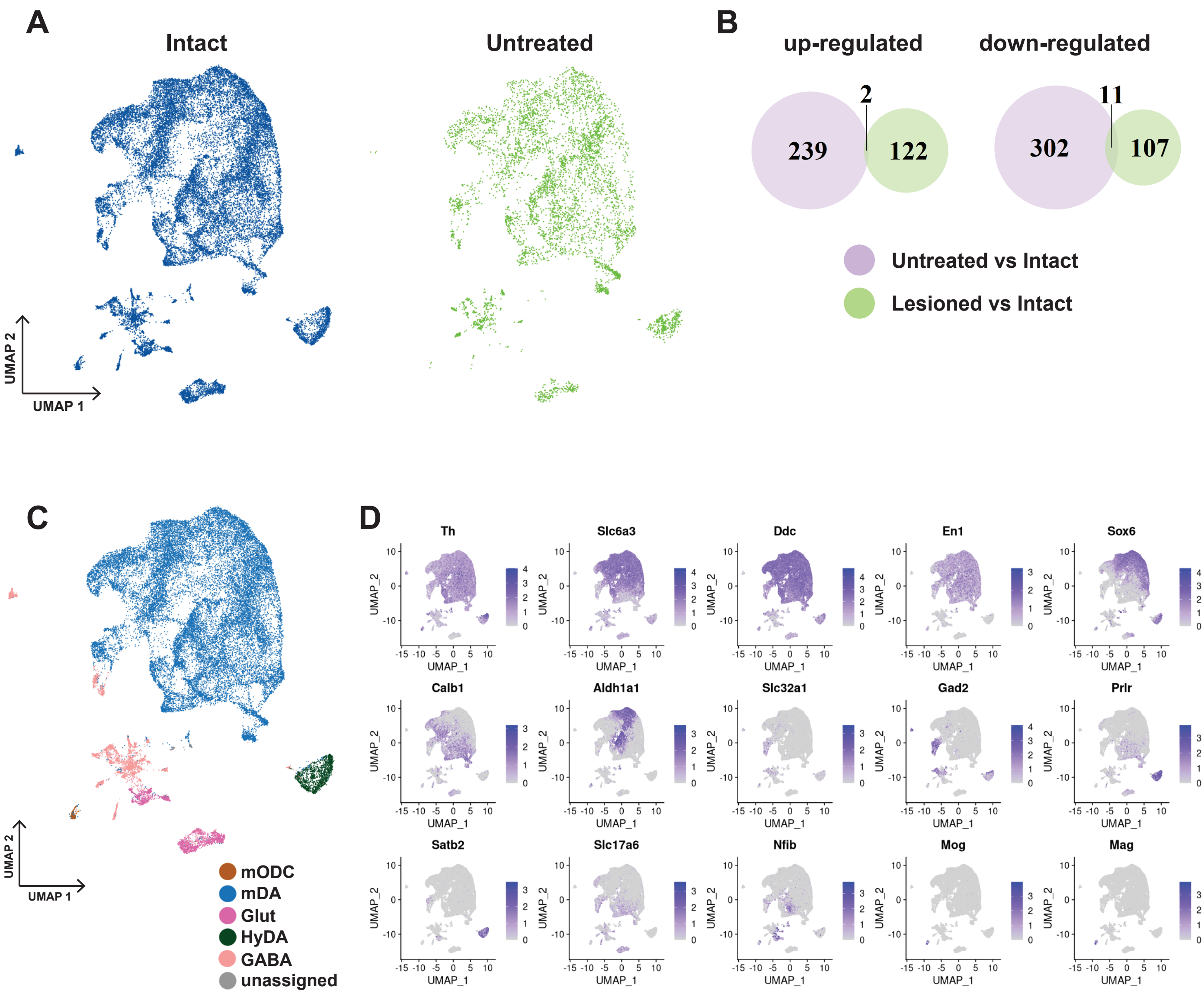

### Figure Supplement 8-1

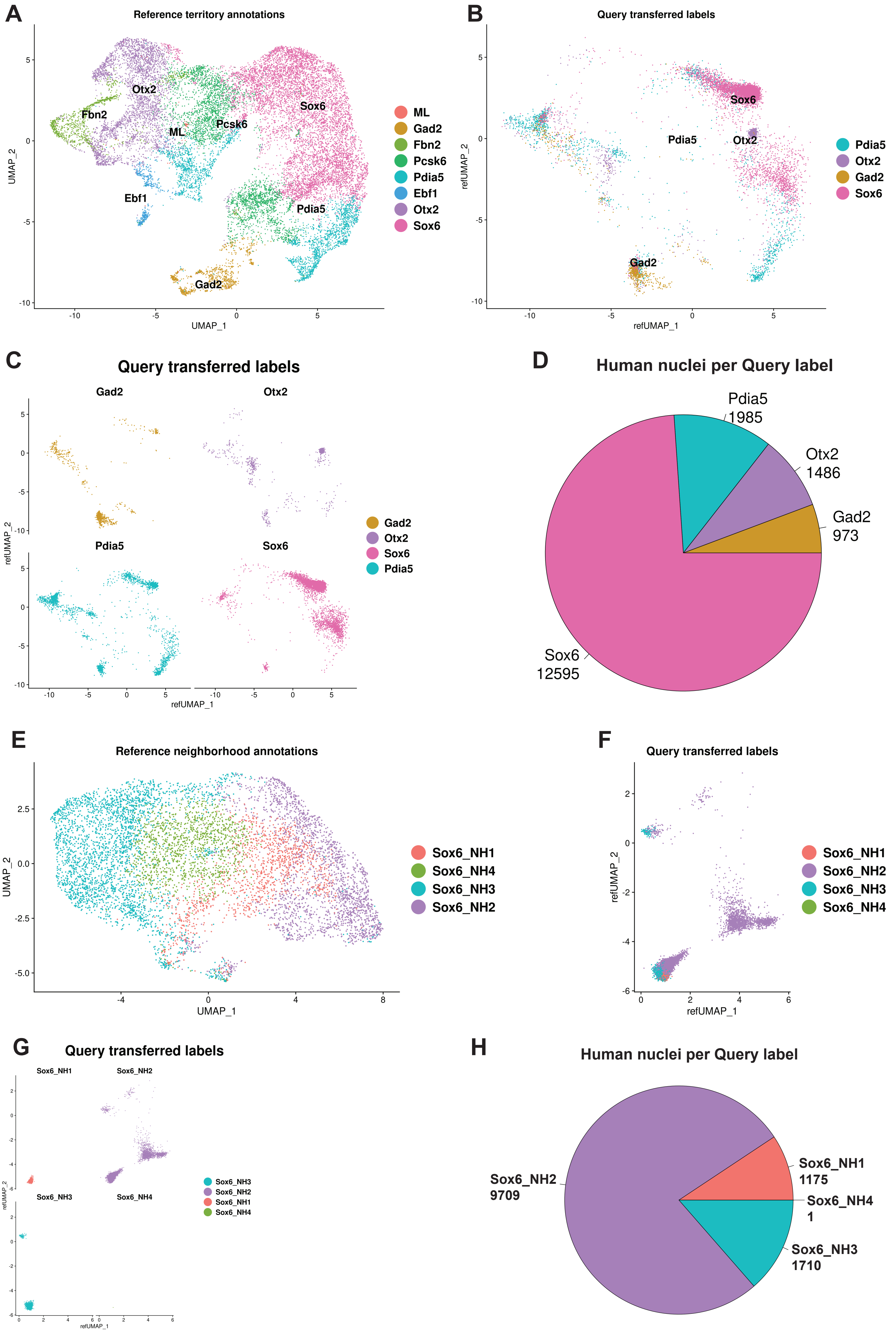
